# Coordination of phage genome degradation versus host genome protection by a bifunctional restriction-modification enzyme visualized by CryoEM

**DOI:** 10.1101/2021.01.06.425610

**Authors:** Betty W. Shen, Joel D. Quispe, Yvette Luyten, Benjamin E. McGough, Richard D. Morgan, Barry L. Stoddard

## Abstract

Restriction enzymes that combine DNA methylation and cleavage activities into a single polypeptide or protein assemblage and that modify just one DNA strand for host protection are capable of more efficient adaptation towards novel target sites. However, they must solve the problem of discrimination between newly replicated and unmodified host sites (needing methylation) and invasive foreign site (needing to lead to cleavage). One solution to this problem might be that the activity that occurs at any given site is dictated by the oligomeric state of the bound enzyme. Methylation requires just a single bound site and is relatively slow, while cleavage requires that multiple unmethylated target sites (often found in incoming, foreign DNA) be brought together into an enzyme-DNA complex to license rapid cleavage. To validate and visualize the basis for such a mechanism, we have determined the catalytic behavior of a bifunctional Type IIL restriction-modification (‘RM’) enzyme (DrdV) and determined its high-resolution structure at several different stages of assembly and coordination with multiple bound DNA targets using CryoEM. The structures demonstrate a mechanism of cleavage by which an initial dimer is formed between two DNA-bound enzyme molecules, positioning the single endonuclease domain from each enzyme against the other’s DNA and requiring further oligomerization through differing protein-protein contacts of additional DNA-bound enzyme molecules to enable cleavage. The analysis explains how endonuclease activity is licensed by the presence of multiple target-containing DNA duplexes and provides a clear view of the assembly through 3D space of a DNA-bound RM enzyme ‘synapse’ that leads to rapid cleavage of foreign DNA.

---

Bacterial restriction-modification (RM) systems are ubiquitous and highly diverse defense mechanisms that guard host cells against invasive DNA elements, particularly phage genomes (Halford, 2009; Loenen et al., 2014b; Roberts, 2005). RM systems pair two competing enzymatic activities: methylation of adenine or cytosine bases within a target site (which protects the host genome from degradation) versus cleavage of DNA within or at some distance from unmethylated copies of the same target site (which leads to degradation of foreign DNA). In combination with additional innate or ‘preprogrammed’ restriction mechanisms that also act on both host and invader genomes (such as the Pgl (Sumby and Smith, 2002), BREX (Goldfarb et al., 2015), DND (Xu et al., 2010) and Ssp (Xiong et al., 2020) defense systems) and complementary ‘adaptive’ nuclease systems (typified by reprogrammable CRISPR-associated nucleases (Koonin and Makarova, 2019)) RM systems represent an important form of antiviral defense in bacteria.

RM systems are loosely divided into at least four major classes, based on their structural composition, biochemical activities and the relationship between their bound DNA targets and subsequent cleavage patterns (Loenen et al., 2014b). Type I and III RM systems contain ATP-dependent translocase domains or subunits that bring together multiple subunits into a DNA-bound protein collision complex or synapse, resulting in cleavage either near (Type III) or at some random distance (Type I) from their target sites (Loenen et al., 2014a; Rao et al., 2014). In contrast, Type II systems do not contain or utilize ATP-dependent motors for motion and activity (Pingoud et al., 2014). Instead, they rely either on the parallel activities of stand-alone methyltransferase (MTase) and endonuclease (Endo) enzymes that independently recognize the same DNA target, or on the physical coupling of methylation and endonuclease domains within a single protein chain or a larger multimeric assemblage, so that both functions are simultaneously targeted by a single DNA recognition module. Type IV endonucleases behave similarly to Type II enzymes, but cleave methylated, rather than unmethylated DNA targets (enabling a bacterial response against phage that methylate their own DNA target to evade restriction endonuclease activity) (Loenen and Raleigh, 2014).

RM systems that use a common DNA recognition module to simultaneously target their competing DNA methylation and cleavage activities have the advantage of facilitating the evolution of new DNA specificity (Morgan and Luyten, 2009), since any alteration in DNA targeting will concurrently alter the specificity of host protective methylation and restrictive cleavage of invading DNAs. Many such RM systems modify just one DNA strand within their asymmetric recognition motif, allowing these systems to employ a single DNA recognition module and MTase domain. This presents a distinct challenge, as DNA replication produces one daughter DNA with no protective methylation. Such systems must therefore solve the problem of discrimination between self (which should be methylated and protected from cleavage) versus non-self (which should be rapidly cleaved and degraded). Systems that communicate between two or more sites within a DNA molecule through 1D translocation, such as the Type III and Type ISP systems, solve the problem by requiring sites be in a head to head orientation to license cleavage, since this effectively places methylation in both strands. However, there are numerous Type IIL systems that do not have a translocase function and cut sites without regard to their orientation. How these avoid self-cutting while maintaining sufficient restriction of invading DNAs to provide a selective advantage to the host has been an open question. One reasonable (and frequently postulated) solution to that challenge is to (1) ensure that cleavage is significantly faster than methylation, while also requiring that (2) multiple unmethylated DNA target sites be brought together into an enzyme-DNA complex before cleavage is licensed to occur. As a result, whereas an encounter with foreign DNA (typically harboring multiple unprotected sites) would lead to rapid cutting at multiple positions, an encounter between an RM enzyme and one or two unmethylated target site(s) on the host would result in eventual DNA methylation and release of bound enzyme.

A variety of structures of RM enzymes that combine their methylation and cleavage domains and activities into single protein chains or complexes have been solved in the presence and absence of bound DNA. These include:

1. Two single-chain Type IIL enzymes (*MmeI* (Callahan et al., 2016) and *BpuSI (Shen et al., 2011),* that each contain an N-terminal nuclease domain followed by methyltransferase (MTase) and target recognition domains (TRDs)).
2. A pair of single-chain Type ISP enzymes (*LlaGI* and *LlaBIII*, that each incorporate an additional RecA-like ATPase domain into their structures (Chand et al., 2015; Šišáková et al., 2013)).
3. A crystal structure for a complex of EcoP15I (a Type III multichain complex containing two MTase subunits and an Endonuclease subunit) bound to DNA (Gupta et al., 2015).
4. CryoEM structures of the Type I enzyme *EcoR124I* (a multichain complex containing multiple nuclease-translocase, methyltransferase and specificity subunits) bound to DNA (Gao et al., 2020). That recent analysis built upon lower-resolution models of DNA-bound ‘M_2_S’ subcomplexes of that same enzyme, as well as those of *EcoKI* and *TteI (Kennaway et al., 2009; Kennaway et al., 2012)*.
5. A Type IV methyl-dependent restriction endonuclease, MspJI, in a tetrameric complex of MspJI bound to DNA (Horton et al., 2012) (although the Type IV do not have a MTase domain, the MspJI tetrameric complex is relevant to this study).

Collectively, these analyses have provided considerable insight into the domain organizations, structural dynamics, DNA recognition specificity, and (for the type ISP *LlaGI* and *LlaBIII* enzymes) a unique mechanism of translocation and subsequent cleavage of DNA (Chand et al., 2015; Šišáková et al., 2013). However, a high-resolution structure of a multimeric RM enzyme system engaged in simultaneous recognition complexes with multiple DNA targets (with the methyltransferase and nuclease domains each properly positioned for competing reaction outcomes) has not yet been described.

DrdV is a single-chain, type IIL restriction-modification enzyme of length 1029 residues that recognizes the asymmetric DNA target site 5’ CATGN**<u>A</u>**C 3’ and methylates an adenine (bold and underlined) in one strand, leading to host protection. It contains an N-terminal nuclease domain, a helical connector region followed by a methyltransferase domain, and a C-terminal target recognition domain (TRD). When bound to its DNA target, it either methylates the underlined adenine within the target or cleaves the top and bottom strand of foreign DNA precisely 10 and 8 basepairs downstream of the target site. In this study, we use CryoEM analysis and supporting biochemical experiments to visualize the stepwise formation of a tetrameric assemblage of DrdV in complex with independently bound DNA target sites. The analysis illustrates the structural basis for generation of an active endonuclease complex of bound enzymes and the basis of crosstalk and cooperativity between multiple copies of the enzyme and bound DNA.

## METHODS

### Protein expression and purification

The gene encoding the DrdV RM system (AXG99744.1) was PCR amplified from *Deinococcus wulumuqiensis* 479 genomic DNA using Q5 hot start high-fidelity DNA polymerase and cloned into the T7 expression-based vector pSAPV6 (Samuelson et al., 2004) using the NEBuilder HiFi DNA Assembly master mix reaction protocol (New England Biolabs, Ipswich, MA). The plasmid construct was confirmed by DNA sequencing of the DrdV gene and flanking vector sequence. The verified plasmid construct was transformed and expressed in the *E. coli* host ER3081 (F^-^ λ-fhuA2 lacZ::T7 gene1 [lon] ompT gal attB::(pCD13-lysY, lacIq) sulA11 R(mcr-73::miniTn10–TetS)2 [dcm] R(zgb-210::Tn10 –TetS) endA1 D(mcrC-mrr)114::IS10).

DrdV endonuclease was purified from 425 g of cells grown at 30°C in Rich media supplemented with 2% glycerol and 0.2% glucose and containing 30μg/ml chloramphenicol. Cells were induced at a final concentration of 0.4mM IPTG and grown for an additional 3 hours at 30°C before harvest. Cells were resuspended in 3 volumes DEAE buffer (300mM NaCl, 50mM Tris pH8, 0.1mM EDTA, 1mM DTT, 5% glycerol), lysed using Microfluidics microfluidizer M110EH (Microfluidics, Westwood, MA) and cell debris removed by centrifugation at 15,000xg for 40min.

DrdV endonuclease was purified to near homogeneity via four sequential chromatographic steps: DEAE anion exchange, Heparin HyperD, Source 15Q, and Source 15S (**Supplemental Figure S1**, **panel a**). The clarified lysate (1375 mL) was first applied to DEAE (200 mL column bed volume, pH 8.0) and then washed with 2 column volumes (400 mL) of DEAE buffer. The flow-through and wash were pooled (2145 mL), diluted with no-salt DEAE buffer (50 mM Tris pH8, 0.1 mM EDTA, 1 mM DTT, 5% glycerol) to a final NaCl concentration of 150 mM, and applied to a heparin HyperD column (pH 8.0). A salt gradient was run from 150 mM to 1000 mM NaCl and 25 mL fractions were collected. DrdV eluted across fractions 40 to 50 (275 mL total volume). Those fractions were diluted to 55 mM NaCl and applied to a SourceQ column, and the protein eluted via a salt gradient while collecting 22 mL fractions. DrdV eluted across fractions 17 to 22 (132 mL total volume; 2700 mg total protein). The fractions were pooled, diluted to 50 mM NaCl and applied to a Source S column at pH 8.0, and eluted via a salt gradient into 20 mL fractions. DrdV eluted across fractions 13 to 22 (120 mL total volume). They were pooled and dialyzed into storage buffer (250mM NaCl, 10mM Tris pH 9.0, 0.1 mM EDTA, 1 mM DTT, 50% glycerol). 1200 milligrams (1.2 grams) of purified protein was stored at a final concentration of 20 mg/mL.

Analytical size exclusion chromatography (SEC) demonstrated that the purified DrdV protein eluted at a volume corresponding to an approximate molecular weight of approximately 100 kilodaltons, corresponding to a monomer in solution. Upon incubation, in the presence of calcium, with an equimolar amount of a double stranded DNA (dsDNA) duplex containing a single enzyme target site (consisting of a top stranded with sequence 5’ -CAGCC*<u>CATGGAC</u>*CCAGAACCAC/CCACC-3’ (<u>underline</u> = target site; “/” = cut site) and its complementary bottom strand with sequence 3’-GTCGG*<u>GTACCTG</u>*GGTCTTGG/TGGGTGG 5’), the protein co-eluted with the DNA at a volume corresponding to an approximate molecular weight of 540 kilodaltons, suggesting the formation of a tetrameric enzyme-DNA complex (**Supplemental Figure S1, panel b**).

## DrdV endonuclease and methyltransferase assays

Endonuclease activity was assayed in NEBuffer 4 (20mM Tris-acetate, pH7.9, 10mM magnesium acetate, 50 mM potassium acetate, 1 mM DTT) supplemented with 80 μM S-adenosyl-methionine (AdoMet), typically using 1 μg DNA substrate per 50 μl reaction volume at 37°C. Reactions were terminated by adding stop solution containing 0.08% SDS (NEB Gel Loading Dye, Purple) and DNA fragments were analyzed by electrophoresis in agarose gels. Methyltransferase activity was assayed in the same buffer, supplemented with 12.5 mM EDTA (to remove Mg^++^) and 80 μM AdoMet.

Cleavage assays that illustrated the *trans-*activation of the endonuclease via the addition of dsDNA harboring the enzyme’s target site were performed in the presence of an added oligonucleotide (sequence 5’-GTGCTCAG<u>GTCCATG</u>AGCGAGTCTTTTGACTCGCT<u>CATGGAC</u>CTGAGCACTC-3’) that forms a short hairpin double-stranded DNA duplex (**Figure 1)**containing the CATGNAC recognition site (top and bottom strands of the target corresponding to the underlined bases in the sequence shown) with 8 basepairs upstream (5’) an 8 basepairs downstream (3’) of the target, terminating immediately prior to the site of DNA cutting.

**Figure 1.**
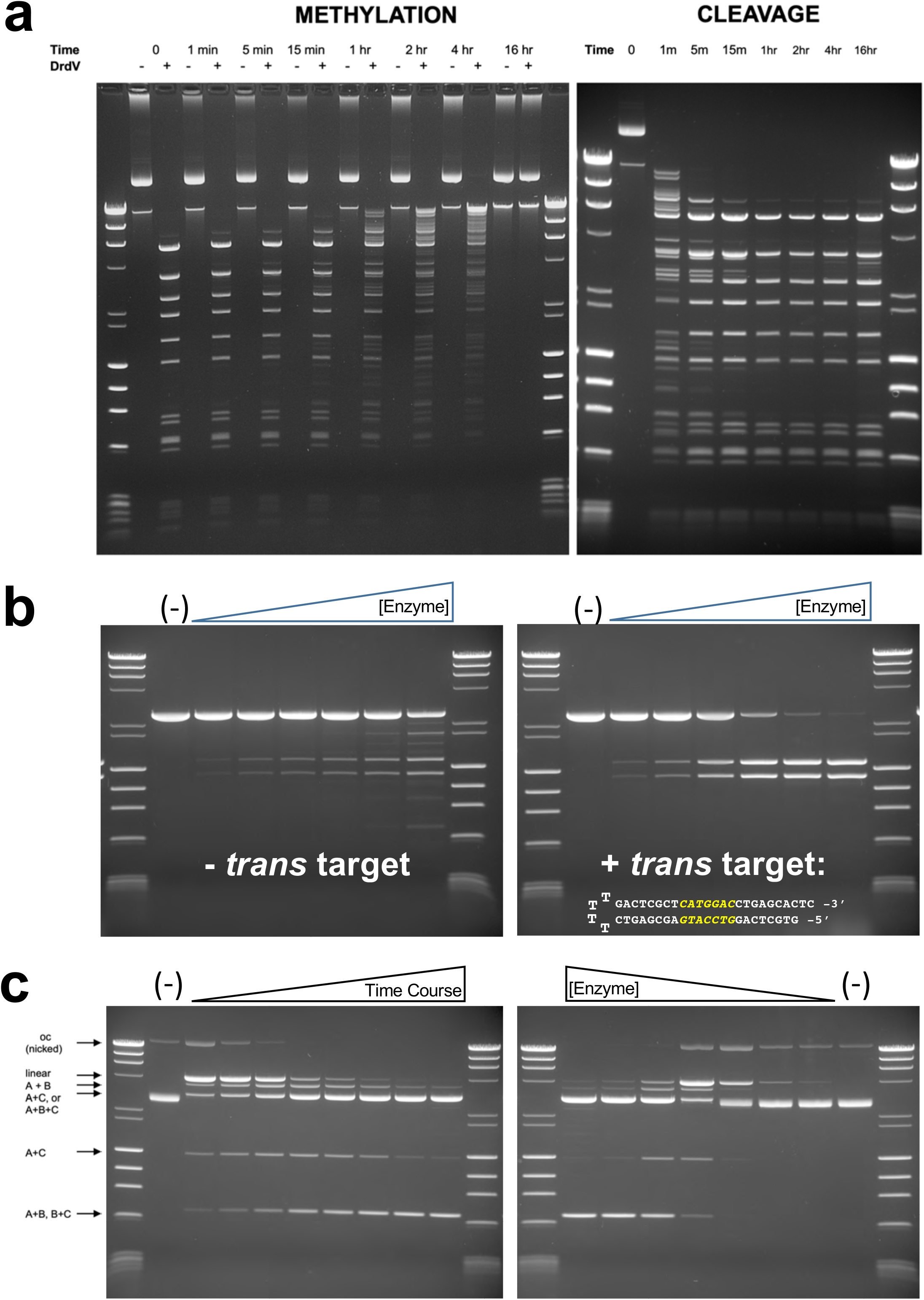
*In vitro* biochemical analyses of DrdV activities. See ***Methods*** for details of all reaction conditions. ***Panel a:*** Methylation by DrdV is much slower than endonuclease cleavage. *<u>Left</u>*: Time course of DrdV incubation in buffer with SAM (AdoMet) but without Mg^++^ (to prevent cleavage). The DNA substrate (pAd2BsaBI plasmid containing 19 DrdV sites) was incubated with 1 unit DrdV for the indicated time, then immediately purified using a spin column. The purified DNA was then challenged by cutting with DrdV (now in the presence of Mg^++^) to assess methylation status. Some partial methylation is observed starting at 15 minutes, but full methylation requires between 4 and 16 hrs. *<u>Right</u>:* Same time course in buffer containing Mg^++^. Cleavage is 90% complete within 5 minutes and fully complete in 1 hour. ***Panel b***: Cleavage is activated by presence of DNA target site added in *trans. <u>Left</u>*: DrdV cleavage of a pUC19 plasmid substrate, (linearized with PstI) that harbors a single DrdV site: 2-fold serial dilution of DrdV from 8 to 0.25 units. The extra bands indicate star activity at near-cognate DrdV sites in the presence of the highest amounts (8 and 4 units) of DrdV. *<u>Right</u>*: DrdV cleavage of the same substrate, at the same enzyme concentrations, each in the presence of 100 mM of a short DNA hairpin duplex containing the DrdV target site. Cleavage goes to completion and displays greatly reduced off-target cutting. ***Panel c***: DrdV cleaves both DNA strands at multiple target sites in a coordinated manner. <u>***Left***</u>: Time course of DrdV digestion of supercoiled pBR322 DNA (2 units/ug) for 15 s, 30 s, 1, 3, 5, 10, 30 and 60 min. Supercoiled plasmid is converted directly to linear plasmid cut at one site, or to fragments representing cutting at two sites, with very little appearance of open circle (OC) DNA nicked in one strand only. Subsequently the DNA is cut at all three sites. *<u>Right</u>*: 2-fold serial dilution of DrdV from 8 to 0.125 units/ug pBR322 substrate. At limiting enzyme, the majority of the cut DNA represents cutting at one site to linearize the plasmid.

### Structural visualization via electron microscopy

The protein-DNA complexes were initially evaluated by negative-stained TEM (**Supplemental Figure S1, panels c, d, e)** followed by screening cooling and vitrification conditions and initial data collection using a GLACIOS 200kV electron microscope (**Supplemental Figure S2**). A subsequent data set was collected on a KRIOS electron microscope (**Supplemental Figure S3**).

All data preprocessing, which include motion correction, ctf estimation, and exposure curation, as well as 2D particle curations, 3D model generation/refinement, and post refinement were performed using the software package cryoSPARC (Punjani et al., 2017). For each movie stack, the frames were aligned for beam-induced motion correction using Patch-motion-correction. Patch-CTF was used to determine the contrast transfer function parameters. Bad movies were eliminated based on a CTF-fit resolution cut off at 5Å and relative ice thickness of 1.2 estimated from the CTF function by cryoSPARC2. Different particle picking algorithms, including manual pick, template-based and blob picking were employed to the same dataset and results on model distribution were compared. The evaluation of the density map at all stages and initial fitting of the Phyre2(Kelley et al., 2015) predicted model to the final density map were accomplished in Chimera (Pettersen et al., 2004). The final structures were built and refined with program COOT (Emsley et al., 2010).

#### I. Negative stain transmission electron microscopy (TEM)

Negative-stain grids (**Supplemental Figure S1, panel c)** were prepared by the application of 4 μl of SEC purified samples to a glow discharged uniform carbon film coated grid. The particles were allowed to adsorb to the surface for 30 to 60 seconds. Excess solution was wicked away by briefly touching the edge of a filter paper. The grid was quickly washed three times with 20 μl drops of water and once with a drop of 20 μl 0.5% uranyl formate (UF) followed by staining for ~20 second with a 40 μl UF. The grids were air-dried for at least 2 hours prior to inspection on an in-house JEOL JM1400 microscope (operating at 120 kV) equipped with a Gatan Rio 4kx4k CMOS detector.

Both DNA free DrdV and DrdV/DNA complex distributed homogeneously in random orientations over the surface of the carbon film. A small dataset of 126 micrographs was collected using the automated data collection package Leginon (Suloway et al., 2005) from the negative-stained specimen at a pixel size of 1.6Å on a FEI Tecnai Spirit electron microscope (operating at 120 kV) equipped with a Gatan 4k × 4k CCD detector.

Initially 2800 particles were hand-picked from all 126 micrographs of the negative stained dataset and subjected to reference free 2D classification. Six out of ten of 2D class averages (**Supplemental Figure S1, panel d)** from 2D classification were used to reconstruct a four-loped volume with imperfect two-fold symmetry. Homogenous refinement with C1 or C2 symmetry yield envelopes at approximately 17 Å and 14 Å resolution at a Gold Standard Fourier Shell Correlation (GSFSC) of 0.143, respectively. **(Supplemental Figure S1, panel e**).

#### II. Initial CryoEM Screening and analyses (see **Supplemental Figure S2**)

Using the same protein-DNA complex, CryoEM grids were prepared by applying 3 μL of a DrdV-DNA complex with an absorbance of 0.55 OD at 280 nm (approximately 0.25 mg/ml protein based on quantitative SDS-PAGE analysis) to a glow-discharged Quantifoil1.2/1.3 holey carbon film coated copper grid, which was blotted for 5.0 s and plunge-frozen in liquid ethane using an FEI Vitrobot Mark IV. Screening datasets with a total of 1410 movies were collected from two separate grids on a GLACIOS electron-microscope (operating at 200 kV) equipped with a Gatan K2-Summit direct electron detector at a pixel size of 1.16 Å. The same six selected 2D classes of the negative stained particles (**Supplemental Figure S1, panel d**) were used as templates for automatic templated particle picking from a total of 1220 movies after frames were aligned and manual exposure curation.

After “inspect particle picks” and “local motion correction”, 354364 out of 703210 particles were accepted for 2D classification. After 3 rounds of particle curation, 82768 particles from 26 selected classes (out of 100) were used for *ab-initio* reconstruction of one unique 3D model. This initial 3D cryoEM reconstruction of DrdV-DNA complex showed an asymmetric particle with three rather than four lobes as shown for the negative-stained particles. which ruled out higher symmetry of C2. Hence all further refinement processes were performed with C1 symmetry to avoid biased interpretation of the resulting maps, resulting with a trimer map of 3.25Å at a GSFSC of 0.143 between the two half maps. 3D variability analysis (Punjani and Fleet, 2020) of this map showed that the most prominent component arises from the association/dissociation of a third protomer to a dimer core (**Supplemental Movie 1**). *Ab-initio* 3D reconstruction with four models revealed the presence of a dimer, a partial trimer with an ill-defined dimer core, and a full-trimer (at 28.4%, 23.8% and 41.8%, respectively) plus a small percentage of smaller fragments (6.0%). Homogeneous, nonuniform refinement followed by Local refinement resulted in a map at 3.3Å for a dimer and a map at 3.4Å for a full trimer (**Supplemental Figure S2** and **Figure 2, panel a**).

**Figure 2.**
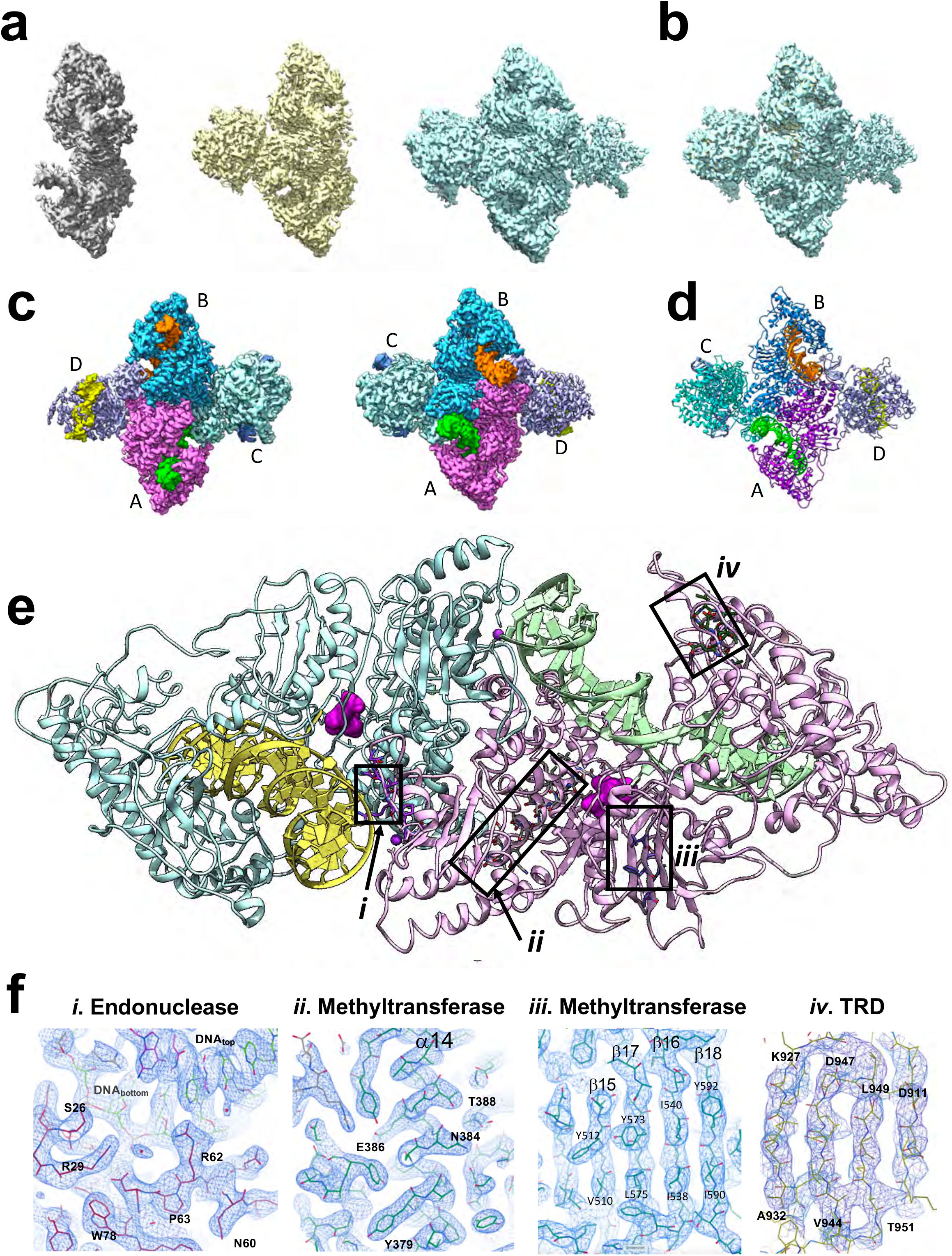
CryoEM analysis of DrdV-DNA complex. ***Panel a***: Density maps of dimeric, trimeric and tetrameric DNA-bound enzyme complexes. The fourth and final subunit in the tetrameric enzyme assemblage is displays sub-stoichiometric partial occupancy. See also **Supplemental Movie 1.***Panel b*: Superposition of all three density maps. ***Panel c:*** Front and back views of the DrdV tetramer density maps with individually colored enzyme subunits and bound DNA duplexes. ***Panel d:*** Atomic model of the DrdV tetrameric assembly. ***Panel e:*** Model of the central DNA-bound enzyme dimer (subunits A and B) extracted from the tetrameric assemblage, overlayed with boxes corresponding to representative regions of density and respective models shown in ***Panel f. Panel f:*** Close-up views of CryoEM map corresponding to (i) the N-terminal endonuclease domain; (ii and iii) the central methyltransferase domain and (iv) the C-terminal target recognition domain (TRD) as indicated by boxes in ***Panel e.***

#### III. Final CryoEM data collection (see **Supplemental Figure S3**)

A final dataset was collected at the Pacific Northwest Center for CryoEM (PNCC) using a vitrified grid prepared with the complex at a final concentration of ~0.4 mg/ml protein (diluted immediately before application to the grid from a stock solution at ~1.6 mg/ml protein) on a Quantifoil1.2/1.3, 200 mesh copper grid, using a Titan KRIOS electron microscope (FEI) operating at 300 kV, equipped with a Gatan K3 direct electron detector and an energy filter (operated with a slit width of 20 eV) at a super resolution pixel size of 0.5318Å. The data was binned by a factor of two to a pixel size of 1.064Å. After preprocessing (motion correction, ctf estimate and manual exposure curation), 3927 micrographs were accepted from a total of 4300 movies. An automated ‘Blob Picker’ algorithm with maximum and minimum diameters of 240 Å and 110 Å was used for particle picking. After inspection and three rounds of particle curation, 376599 particles were selected for 3D reconstruction and refinements. *Ab-initio* 3D reconstruction with three models showed that the dataset contained three different classes – trimers (~34%), tetramers (~50%), and a higher molecular aggregate (~15%) that could be the result of the addition of extra monomers to the tetramers or the result of close contact of neighboring particles. Dimers were absent in the PNCC dataset which was prepared from a stock solution at much higher concentration and diluted immediately prior to the preparation of the grids.

After homogeneous and non-uniform refinement, the refined tetramer class was further refined after Local- and Global-CTF refinement of the particles, which led to a final resolution of 2.73 Å at a GSFSC of 0.143 between the two half maps. 3D variability display showed that the association-disassociation of a fourth component to the trimer is the most prominent contribution to the 3D variability, thus the particles under the tetramer class (**Supplemental Figure S3**, tetramer I) were further classified via a second round of *ab-initio* 3D reconstruction with three models, resulting in a full trimer (45.2%), a tetramer (50.2 %) and a class of small fragments (4.6%). Final refinements of the trimer and tetramer led to a resolution of better than 2.9Å at a GSFSC of 0.143 for both 3D classes (**Supplemental Figure S3**). Even though the nominal resolution of Tetramer I is higher than Tetramer II, the quality of the latter is actually superior, especially the Mtase and TRD domains and DNA of the fourth protomer.

The density maps of all the enzyme-DNA complexes, visualized at three unique stages of assembly (dimers, trimers and full tetramers) each allowed unambiguous placement of individual protein chains, each containing all 1029 residues with the exception of a short surface exposure loop in the methyltransferase domain (residue 412 – 421) (**Supplemental Figure S4**), as well as bound copies of the DNA duplex, a bound SAM cofactor, a base-flipped adenine nucleotide in each methyltransferase active site, and calcium ions associated with each endonuclease domain in contact with a DNA strand and scissile phosphate. The map of the tetramers indicated a reduced occupancy for one of the subunits and its bound DNA duplex.

## Results and Discussion

### Biochemical activity assays

A series of *in vitro* biochemical analyses (**Figure 1)** of DrdV activity demonstrate that the DrdV enzyme displays mechanistic behavior described above, that is believed to lead to different reaction outcomes against ‘self’ versus ‘foreign’ DNA targets: much slower methylation than cleavage, strong activation of cleavage via binding of multiple DNA targets, and coordinated, near-simultaneous cleavage of both strands in multiple target sites within a DNA substrate.

#### (i) Cleavage of unmethylated targets by DrdV is significantly faster than the rate of host-protective methylation (**Figure 1a**)

In a series of in vitro incubations with a standard multisite substrate (lambda DNA), DNA cleavage is nearly complete within 1 to 5 minutes, whereas complete methylation of the same substrate under similar conditions (except for the absence of Mg^++^ to prevent DNA cleavage) requires up to 16 hours.

#### (ii) DrdV requires multiple sites for efficient, high fidelity cleavage

DrdV cleaves a DNA substrate containing a single target site (a pUC19 plasmid with a DrdV target site added at position 1680) incompletely, cutting only around 20% of the DNA even with an 8-fold excess of enzyme (**Figure 1b. left**). At higher excess enzyme, star activity (cleavage at closely related non-cognate sites) appears as DrdV begins to make additional, though very partial, cuts at near-cognate sites. In contrast, cleavage activity towards the same plasmid substrate is significantly increased (and off-target star activity is reduced) by supplying a short dsDNA hairpin oligonucleotide that contains the DrdV recognition site *in trans* (**Figure 1b, right)**. Maximum stimulation is achieved when the oligo is supplied at a ratio of between 1:1 to approximately 6:1 to DrdV enzyme molecules, with enzyme molecules in excess to the substrate target sites to be cut.

#### (iii) DrdV simultaneously cleaves both DNA strands, downstream of the target site, in a coordinated manner

DrdV digestion of a circular DNA substrate (pBR322 plasmid) containing multiple target sites showed little accumulation of the nicked open circular DNA form (**Figure 1c)**, indicating that cleavage events occur in a coordinated reaction with both strands cleaved in a nearly concerted event. The plasmids were initially cut both to linear fragments cut at one site only and to fragments cut at two sites, indicating cleavage can occur at just one site or at two sites in a coordinated manner.

### CryoEM Structural analysis

The purified enzyme eluted with an apparent mass of 118 kD from a final size exclusion column. Upon incubation with an equimolar ratio of a DNA duplex containing a single copy of its DNA target site, the protein co-eluted with the DNA over a sharp peak centered at an estimated molecular weight of approximately 540 kD. That sample was used for negative stain EM studies and single particle reconstruction, resulting in a molecular envelope corresponding to an asymmetric tetrameric assemblage with a pair of pseudo-orthogonal dyad symmetry axes, of approximate size 200 × 220 × 100 Å (**Supplemental Figure S1*e*)**. In addition to these largest particles, smaller particles corresponding to intermediate bi- and tri-lobed assemblages were also observed. We interpreted this result as potentially representing a population of enzyme tetramers bound to multiple DNA targets, interspersed with smaller dimeric and trimeric enzyme-DNA assemblages.

The subsequent CryoEM single particle reconstructions showed that DrdV-DNA complexes undergo a concentration dependent oligomerization producing density maps with two, three and four lopes corresponding to dimer, trimer and tetramer and allowed unambiguous placement and subsequent building and refinement of unique enzyme-DNA complexes containing two, three or four protein subunits. All particles contain a highly homologous dimeric core with one or two extra protomers at either side of the trimer and tetramer (**Figure 2, Supplemental Figures S2 and S3, Supplemental Movies**). The final maps (individually corresponding to 3.5 to 2.8 Å resolution) provided well-resolved features that allowed unambiguous modeling of secondary structure elements and corresponding side chain positions across four sequential functional regions and folded domains within each protein subunit (an N-terminal endonuclease domain, an alpha-helical connector, a methyltransferase domain and a C-terminal target recognition domain, or ‘TRD’) of the enzyme (Figure 2, and Supplemental movie 3). The relative local resolution distribution of all three density maps on the same scale and the sequence of a DrdV subunit with corresponding secondary structures are shown in **Supplemental Figure S4**.

The description of the enzyme-DNA complex features provided below is derived from the density map of the largest observed enzyme assemblage. Those points are also observed in the structures of the dimeric and trimeric species (solved and refined independently), with the exception of small conformational changes that appear to accompany the stepwise addition of the third and fourth enzyme subunits.

The relative domain orientations within a single DNA-bound enzyme subunit and its interactions with its target site (observed in all of the structures) is illustrated in **Figure 3**. The DNA target site (numbered according to their position in the target site, i.e. ‘5 - C_1_A_2_T_3_G_4_G_5_<u>A</u>_6_C_7_ - 3’; **Figure 3a**) is bound in a cleft between the methyltransferase domain (MTase, residues 295-635) and target recognition domain (TRD, residues 636-1029) (**Figure 3b)**. The adenine base at position 6 (‘<u>A</u>_6_’) is flipped into the methyltransferase active site and positioned near a bound molecule of S-adenosyl-methionine (‘SAM’ or ‘AdoMet’) (**Figure 3d,e**). Sequence-specific base contacts by enzyme side chains are observed to six (out of the seven) base pairs within the target site, with only the guanine at the fifth position in the target site (where any basepair is tolerated by the enzyme) excluded from direct readout (Figure 3 and Supplemental Figure S5). Individual side chain contacts to the target bases are formed both by the MTase domain (N448, Q485, K486, K488, N548, R554, D564) and by the TRD (N673, R721, Y764, D803, K807). The flipped adenine base is bracketed by π-stacking with F304 and Y451 and forms additional contacts with F562 and N448. The space vacated by the flipped adenine <u>A</u>_6_ is occupied by N548 and K567 from the side of the major groove.

**Figure 3.**
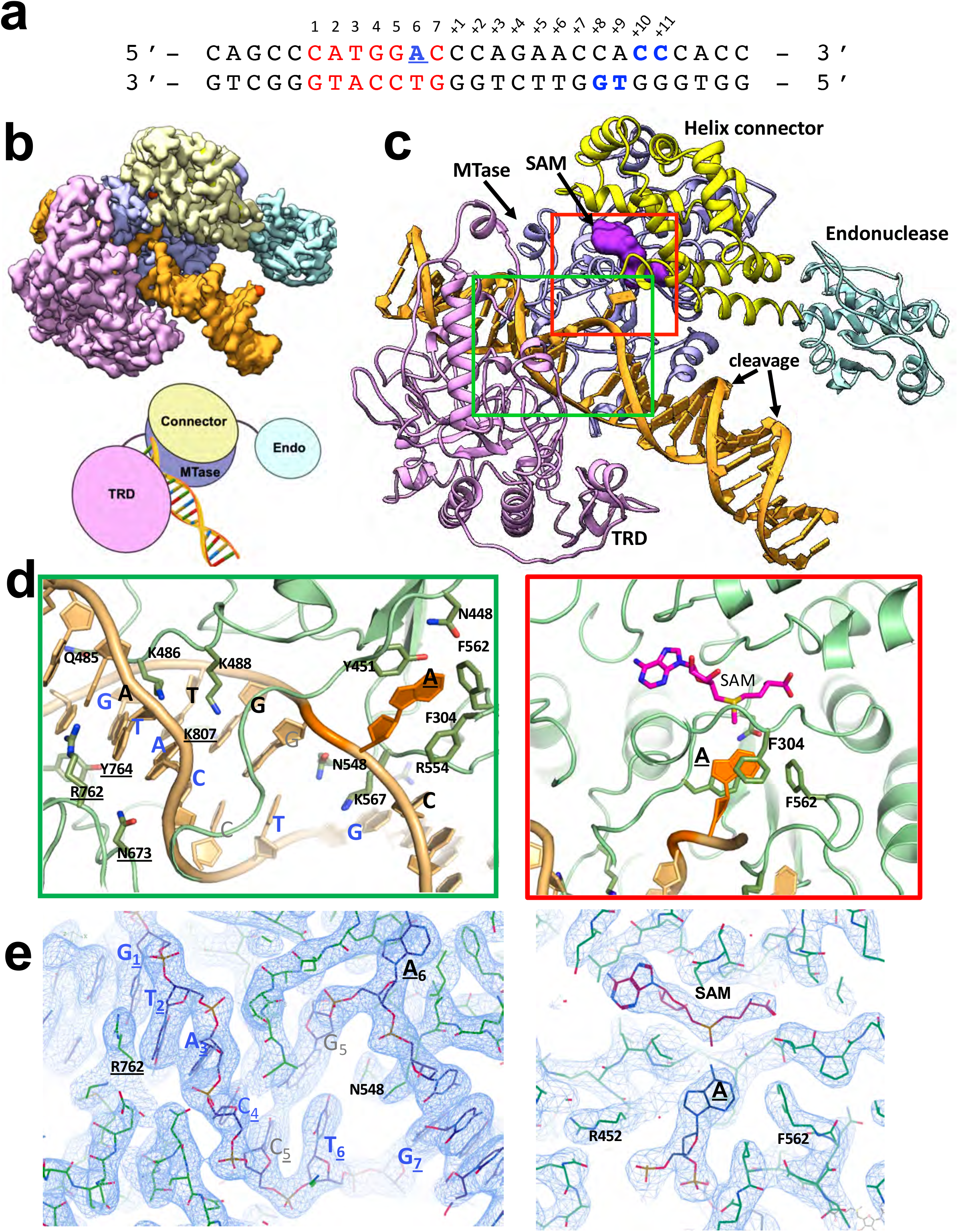
Conformation of an individual DrdV subunit bound to a DNA target. The model and maps shown are extracted from the tetrameric enzyme assemblage. ***Panel a:*** DNA construct used for CryoEM analyses. The duplex consists of 28 complementary basepairs and spans both the enzyme’s seven basepair target site (red bases and blue underlined site of adenine methylation) and its downstream cleavage sites on the top and bottom strands (blue bases flanking the scissile phosphates, 10 and 8 basepairs downstream from the final basepair of the target site). ***Panel b:*** Density map of a single DNA bound DrdV protomer with color coded domains: DrdV is a single chain protein spanning 1029 residues, corresponding to an N-terminal endonuclease domain (light blue), a subsequent helical connector region (yellow) and central methyltransferase (‘MTase’) domain (purple) and a C-terminal target recognition domain (‘TRD’) (pink). ***Panel c:*** Each enzyme subunit contains a bound S-adenosyl-methionine (‘SAM’) cofactor (magenta) bound in its active site. The adenine base at position 6 in the target is flipped into the MTase active site and is unmethylated. The base is contacted by three aromatic residues (Y451, F304 and F562) and a neighboring asparagine (N448) from the MTase domain. N458 and K567 occupy the space vacated by the filliped-out adenine. ***Panels d and e:*** Views of molecular model and corresponding electron density map in the enzyme-DNA interface, with several residues that form additional sequence-specific contacts to the DNA target site shown. Several basic and polar residues from the methyltransferase (including K486 and K488) and the TRD (including D803 and K807) contribute additional base-specific contacts in the target site. The adenine that is targeted for methylation (underlined ‘<u>A</u>’) is clearly flipped out of the DNA duplex and positioned proximal to the bound SAM cofactor; both moieties are clearly visible in the CryoEM density maps. Additional details of basepair-specific contacts are illustrated in **Supplemental Figure S5.**

Within the DNA-enzyme complex formed by a single DrdV subunit as described above, the endonuclease domain is not in contact with the bound DNA duplex (**Figure 3bc**). Instead, it contacts the top strand and the corresponding scissile phosphate in the DNA duplex bound by the opposing subunit within a central dimeric enzyme assemblage (**Figure 4ab**). The endonuclease domain from the opposing enzyme subunit is similarly domain swapped; both endonuclease domains are properly positioned to cleave the top strand of the DNA duplex that is engaged by the opposing enzyme subunit (**Figure 4c**).

**Figure 4.**
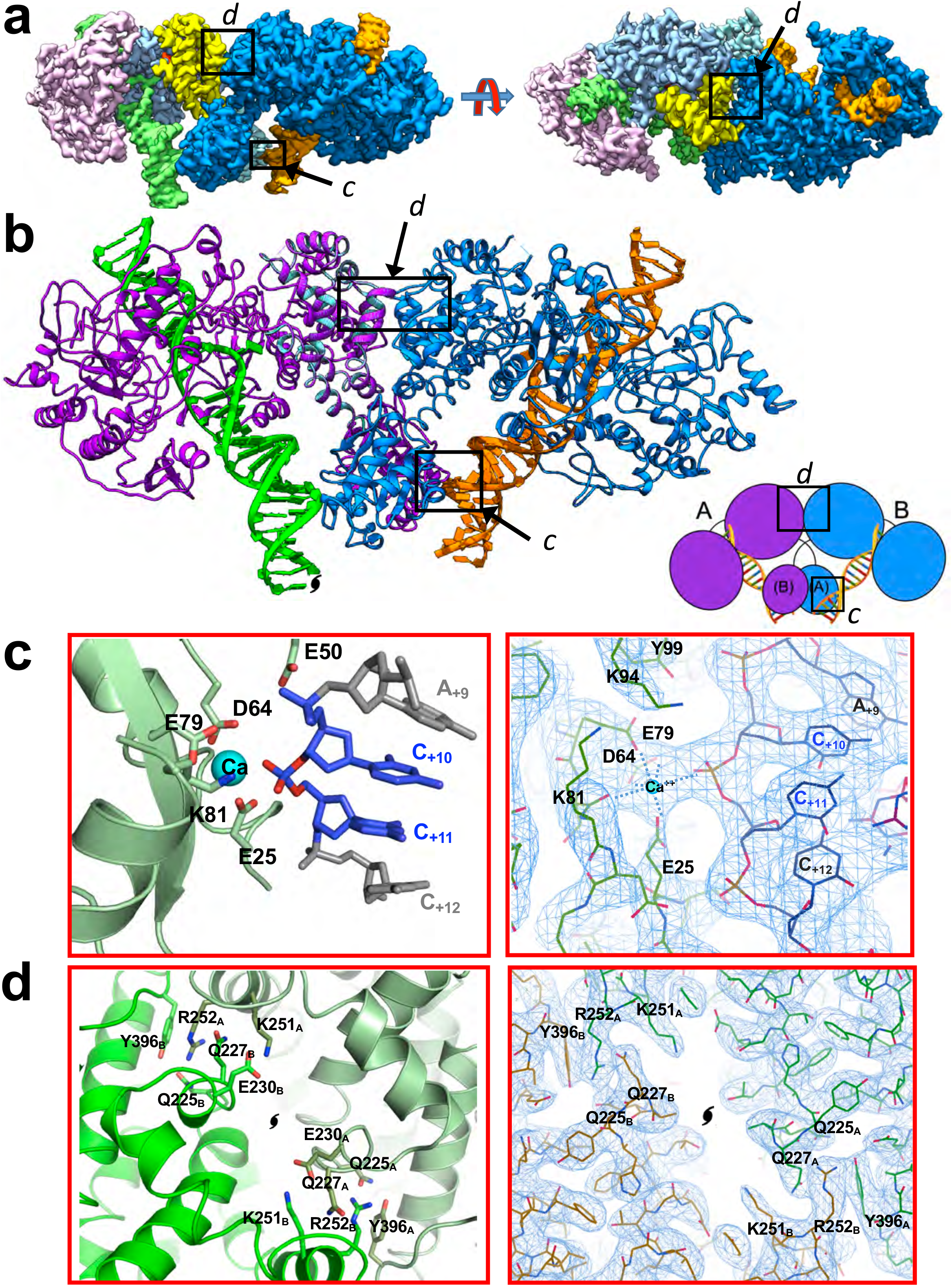
Formation of a DrdV dimer bound to two individual DNA targets positions their endonuclease domains near a one strand of their partner’s bound DNA duplex. The map shown is extracted from the larger tetrameric assemblage. *Panel a:* Two different views of the map. The map corresponding to one enzyme subunit is colored solid blue, while the second is colored to indicate the enzyme’s individual domains. The primary interface between individual protein subunits, and the interface between the DNA duplex bound by subunit B and the endonuclease domain of subunit A, are indicated by boxes labeled ‘*c*’ and ‘*d*’ and correspond to further detail illustrated in *Panels c and d* below. *Panel b:* Ribbon diagram of the DrdV core dimer, again indicating the location of interfaces between the protein subunits (largely via their helical connector regions) and between the nuclease domain of subunit A with DNA target from subunit B. As illustrated in the adjacent cartoon, the endonuclease domains are swapped between subunits, such that each enzyme subunit positions its endonuclease domain in contact with the DNA duplex bound by the opposite enzyme subunit. *Panel c:* Contacts between the endonuclease active site of subunit A and the DNA duplex bound by subunit B, Illustrating the coordination of a bound calcium ion by residues of the active site and by the scissile phosphate on the corresponding strand of DNA. *Panel d*: Ribbon model and density illustrating the interface between the enzyme subunits. The interface is largely composed of two buried, symmetry-related clusters of opposing charged residues (3 acidic side chains (D224, E230 and D231) from one subunit and 3 basic residues (K251*, R252* and K259*) from the opposite subunit, and vice-versa), augmented by similarly duplicated cation-pi interactions between R252 from one subunit and Y396* from the other, as well as an additional contact between R252 and Q225*.

The enzyme dimer displays an extensive buried interface between the two helical connector domains, largely composed of a pair of buried, symmetrically equivalent electrostatic networks and surrounding hydrophobic and hydrogen-bonded contacts with neighboring residues (**Figure 4d**). Within each network, a cluster of three acidic residues from one subunit (D224, E229 and D230) is engaged with a corresponding cluster of three basic residues form the opposing subunit (K251, R252 and K259), thereby bringing together at least 12 opposing charged residues. This electrostatic network is augmented by two patches of electrostatic π-stackings between R252 of one subunit and Q27 andY396 of a second subunit and *vice versa*. In fact, Y396 of the MTase domain is the one and only residue outside of the helical connecter domains that is involved in the interactions between the two subunits in the core dimer.

In each endonuclease-DNA interface (**Figure 4c**) the scissile phosphate is engaged in contacts with a divalent metal ion (a calcium, which was present in the enzyme buffer to prevent cleavage) complexed by a pair of conserved acidic residues (D64 and E79). A neighboring lysine residue (K94) and nearby additional glutamic acid (E25) complete the nuclease active site. The conserved lysine of the canonical PD-ExK endonuclease motif (K81, mutation of which abolishes catalysis) is also positioned near the scissile phosphate where it could participate in catalysis upon adopting a different rotamer than that observed with calcium present in the structure.

The dimeric assemblage of enzyme-DNA complexes described above is further augmented by additional bound DrdV subunits, forming DNA-bound trimeric and tetrameric complexes (**Figure 1 and Figure 5**). (In the tetrameric particles, the fourth and final subunit displays partial, sub-stoichiometric occupancy). In those structures, the additional enzyme subunits are positioned on either side of the dimer described above, via an additional protein-protein interface between two endonuclease domains (**Figure 5a)**). This interface is again composed primarily of a pair of symmetry-related clusters of opposing charged residues (**Figure 5b**), each of which corresponds to K8 and D15 from one subunit forming a pair of buried electrostatic contacts with E41 and R46 from the opposing subunit. The additional endonuclease domain also contacts an extension from the TRD domain of the enzyme bound to the DNA strand being cleaved. The dimerization of the two endonuclease domains places a pair of active sites in alignment with the appropriate scissile phosphates of each strand in a DNA duplex, which in turn allows the enzyme to generate a double strand-break (corresponding to a 2-base 3’ product overhang) downstream of the bound target site **Figure 5a box** and **inset**).

**Figure 5.**
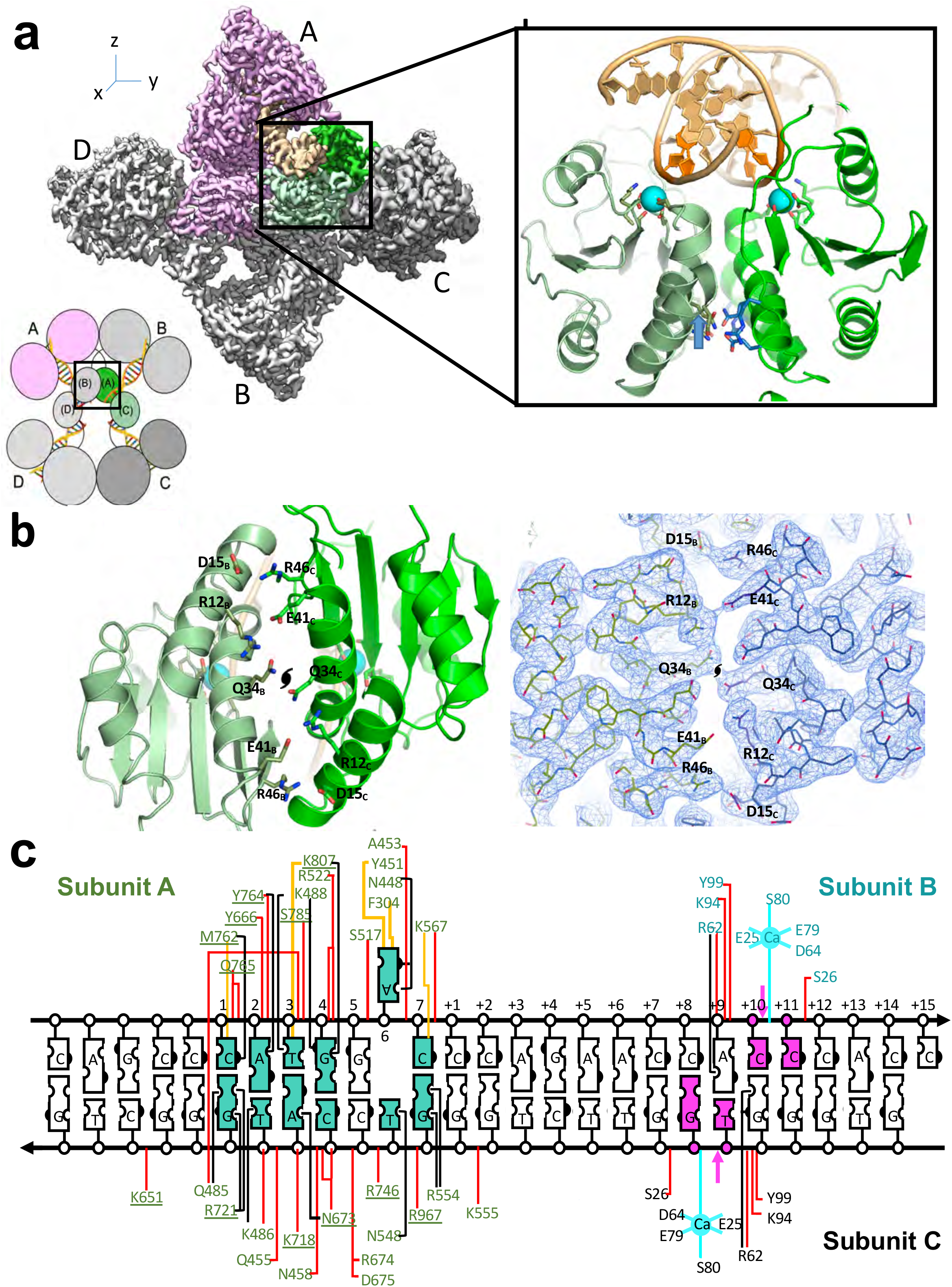
Association of endonuclease domains in the enzyme trimer and tetramer assemblages results in three enzyme subunits being jointly involved in the cleavage of a single bound DNA duplex. ***Panel a:*** Density map of the DNA-bound enzyme tetramer, colored by subunit. The organization of the assemblage and corresponding coloring is also indicated in the cartoon schematic adjacent to the map. The endonuclease domains from subunits B and C (boxed and colored in shades of green) are jointly positioned directly adjacent to the scissile phosphates of the DNA duplex bound by subunit A, with their active sites appropriately arranged to cleave the DNA duplex downstream from the bound target site. The endonuclease domains from subunits A and D are similarly positioned to cleave the DNA duplex bound by subunit B. ***Panel b***: Orthogonal view of the packing and interactions between endonuclease domains shown in ***Panel a,*** and corresponding density map overlayed with atomic model, showing buried, symmetry-related clusters of opposing charged residues (E41 and R46 from one subunit, versus R12 and D15 from the other, and vice-versa) interacting in a pairwise manner within a helical interface (located on the opposite side of the domains from their active site). ***Panel c:*** The recognition, binding and cleavage of a single DNA duplex involves contacts and interfaces formed between three separate DNA-bound enzyme subunits.

As a result of the assembly and coordination of the cleavage complex described above, not less than three individual protein subunits and two bound DNA targets are required in order to form all contacts necessary to cleave a single bound DNA duplex (**Figure 5c**) and four protein subunits are required in order to simultaneously cleave two DNA duplexes.

## Discussion and Conclusions

The biochemical activities and corresponding structures presented in this study reinforce (and illustrate a structural basis for) the concept that an invading DNA such as a phage genome, that presents multiple unmodified sites in a single construct, will be rapidly cleaved whereas the generation of individual unmodified sites in one daughter chromosome following replication would favor modification, as there is less chance to assemble multiple site-bound molecules into cleavage competent complexes. The inefficient cleavage of single site substrates, and activation by specific DNA target sites in *trans,* indicates DrdV must interact with multiple sites to achieve rapid and efficient DNA cleavage.

The structures of DrdV described here (in several stages of assembly with multiple bound DNA targets) offer a sequential view of such an enzyme before and during each stage of DNA search, encounter, coordination and cleavage. When considered alongside previous crystallographic structures of two related Type IIL enzymes at earlier points in their action (BpuSI, solved in the absence of a bound DNA target (Shen et al., 2011) and MmeI, bound to a single copy of its DNA target, in a monomeric complex (Callahan et al., 2016)), a rather complete picture of the functional cycle and mechanism for such bifunctional R-M enzymes seems to emerge. In those earlier crystal structures, the N-terminal endonuclease domain in unbound BpuSI was found to be well-resolved and packed against the interface between its downstream MTase and TRD domains, in a manner that would require its release in order to bind DNA, effectively sequestering the endonuclease catalytic site to prevent any DNA cutting. In contrast, the endonuclease domain in the DNA-bound MmeI enzyme was unobservable (and presumably displaying considerable motion and flexibility), suggesting release from the sequestered position and search for a partner upon initial recognition and engagement of its specific target site.

Like DrdV, MmeI requires multiple sites for cutting and is stimulated by *in trans* DNA containing a recognition site. A simple model of Type IIL enzyme function in a cell would be one in which the apo enzyme is in an inactive (endonuclease-sequestered) state that scans DNA. Upon encounter of its specific recognition motif, the enzyme engages in a tight, long-lived complex that releases the endonuclease domain, but the endonuclease domain is not in contact with the DNA bound by the enzyme. Target recognition and binding would then be followed by a kinetic competition between two outcomes: eventual methylation and enzyme release or encounter and capture of an additional target-bound enzyme subunit to form a dimer with exchange of DNA helices to the partner’s endonuclease domain. Formation of the central dimer is again followed by a kinetic competition between two outcomes: eventual methylation and enzyme release if no additional DNA-bound partners are encountered, or encounter and capture of an additional target-bound enzyme subunit or two to form a catalytically competent trimer or tetramer, leading to rapid cleavage of the two DNAs bound by the central dimer subunits.

The structural analyses presented here also demonstrate that relatively little conformational difference exists between the two core DNA-bound DrdV subunits present in the central dimer particles, and the two additional DNA-bound enzyme subunits that bind to the dimeric assemblage through their endonuclease domains to form the cleavage competent complex. However, examination of differences between those structures does indicate that the formation of endonuclease dimers at each DNA cleavage site (corresponding to the conversion from DNA-bound dimers to larger trimeric and/or tetrameric complexes) is accompanied by observable deformation of the DNA substrates as part of the cleavage mechanism within each bound DNA duplex, and a hinged rigid body rotation of the endonuclease domain by approximately 30° in the catalytic partner subunits relative to that in the central dimer. This rotation highlights the importance of dynamic flexibility of the endonuclease domain relative to the MTase and TRD portion of the protein.

The MTase and DNA recognition domains of DrdV and those of the Type ISP enzymes LlaGI and LlaBIII (Kulkarni et al., 2016) are highly similar, indicating evolution from a common origin, yet the way DNA restrictive cleavage is achieved and controlled between these Type IIL and Type ISP RM systems is quite different. The Type ISP license their endonuclease for cutting through collision encounter between enzymes translocating on the DNA in the opposite direction from inverted recognition sites. Their endonuclease domains never actually encounter one another, but simply nick one strand of the DNA multiples times on either side of the collision complex, eventually leading to double strand breaks when the nicks occur close together as the enzymes move against one another. In stark contrast, DrdV remains bound to its recognition site and recruits additional DNA-bound enzyme molecules, first to form a non-catalytic dimer using one set of protein contacts between their linker and methylase domains that positions the endonuclease of each subunit against the DNA of the other, and then to form catalytic complexes through a different set of protein contacts, largely between endonuclease domains, to bring two endonuclease catalytic centers together for double strand cleavage at a fixed distance from the bound recognition site. This implies the endonuclease domains of Type IIL enzymes are under greater evolutionary pressure and corresponding sequence constraint as they form multiple protein-protein contacts as well as positioning the catalytic center for cutting.

Restriction-modification systems such as those that rely on recognition and cleavage of specific target sequences in foreign DNA are complemented by additional phage restriction mechanisms and systems (such as the Pgl (Sumby and Smith, 2002), BREX (Goldfarb et al., 2015), DND (Xu et al., 2010) and Ssp (Xiong et al., 2020) systems) that also utilize a site-specific protective activity (usually a methyltransferase) to again protect the bacterial genome from self-destruction. The exact manner in which the self-modifying protective activity is employed differs between systems (for example, the methyltransferase activity in the Pgl and BREX systems requires the presence of one or more additional protein factors in order to methylate host DNA). Regardless, the observations described here, which demonstrate the basis of at least one mechanism by which protective versus destructive activities in a restriction system can be biased towards self and foreign, respectively, may be reflected (with many possible variations on a theme) within a wide range of alternative forms of cellular defense.

## Acknowledgements

This work was supported by NIH grant R01 GM105691 to BLS, by an Amazon Cloud Credit to BWS, by the Fred Hutchinson Cancer Research Center, and by New England Biolabs. A portion of this research was supported by NIH grant U24GM129547 and performed at the PNCC at OHSU and accessed through EMSL (grid.436923.9), a DOE Office of Science User Facility sponsored by the Office of Biological and Environmental Research. We thank Justin Kollman and David Veesler at the University of Washington for advice and assistance, Janette Myer for Krios data collection, Jeff Tucker and Dan Tenenbaum for assistance in AWS EC2 setup, Melody Campbell for critical reading of the manuscript, and Sue Biggins and Richard Roberts for support, encouragement and advice.

---

**Supplemental Figure S1.**
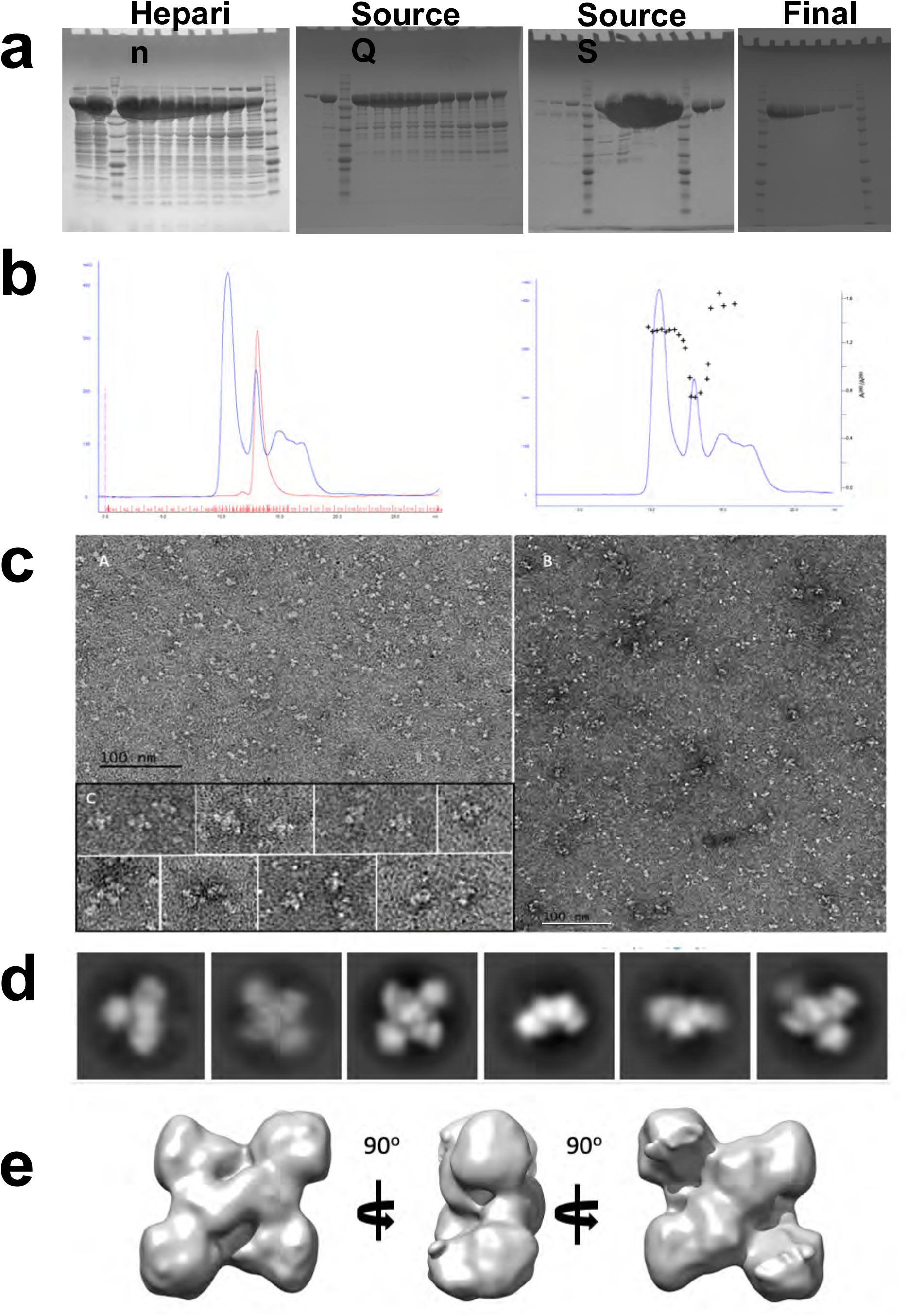
Enzyme purification and initial negative stain EM microscopy. *Panel a.* SDS-PAGE of DrdV at different stages of purification. See methods for full details of enzyme production *Panel b.* SEC elution profiles of free and DNA bound DrdV, red and blue curve respectively (left) and elution profile of DNA bound DrdV overlayed with absorbance ratio at 260 nm and 280 nm, indicating formation and elution of DNA-bound enzyme complex. *Panel c.* Negative stain electron microscopy of DrdV apoenzyme and DNA-bound complex. *<u>A</u>:* DrdV in the absence of DNA. *<u>B</u>:* Negative-stained image of DrdV in the presence of an equimolar amount of DNA duplex (sequence provided in Methods) containing the enzyme target recognition site (CATGGAC) and 16 basepairs downstream of the target. *<u>C</u>:* Selected panels of the DrdV DNA complex. *Panel d:* Selected 2D particle classes., *Panel e*: reconstructed low-resolution 3D model of negative-stained DrdV-DNA particles.

**Supplemental Figure S2.**
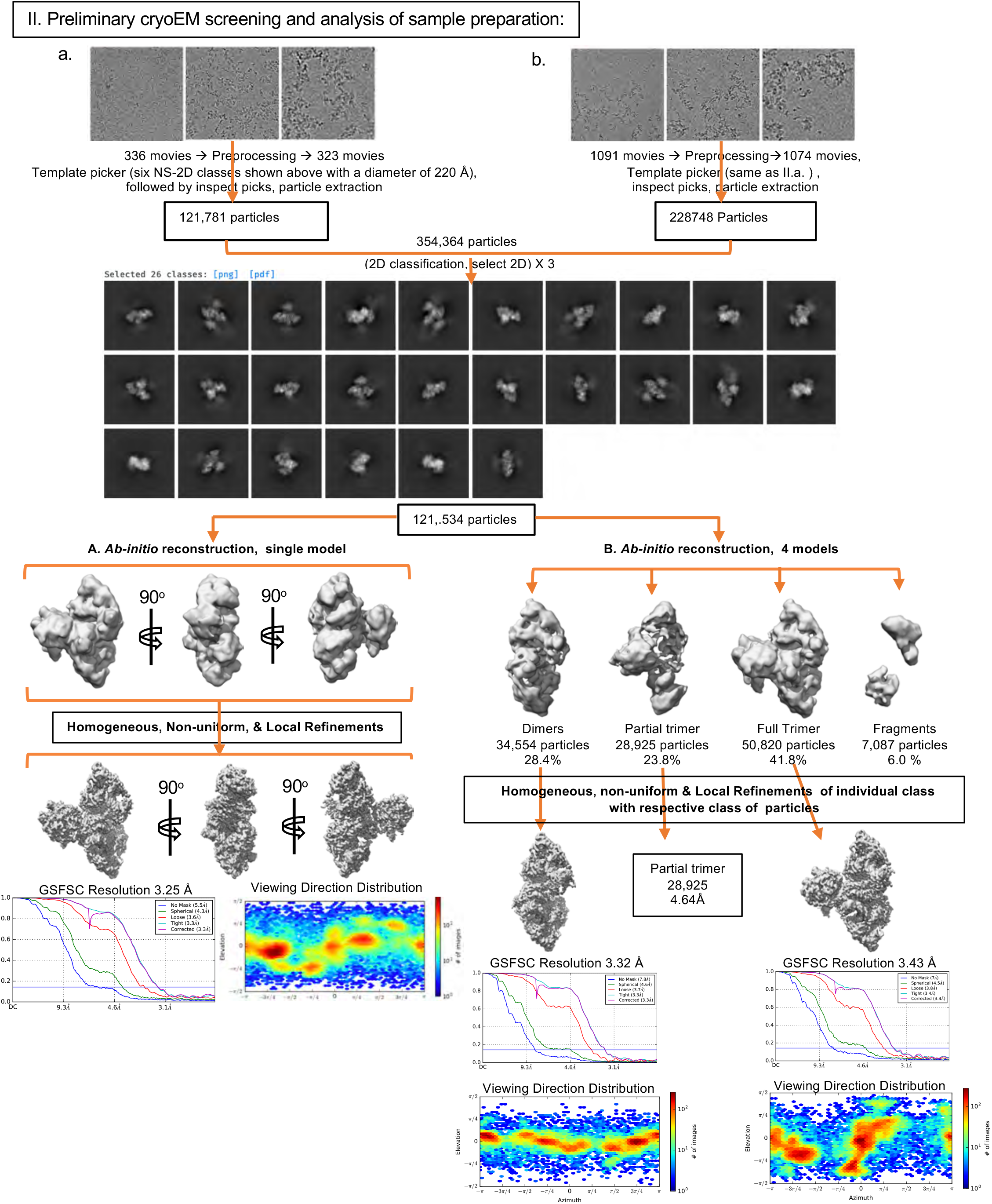
Flow chart for CryoEM analyses using a GLACIOS microscope operating at 200 kV. Data was collected a pixel size of 1.16Å and processed using the package cryoSPARCv2. For full details of data collection and processing approach, see Methods.

**Supplemental Figure S3.**
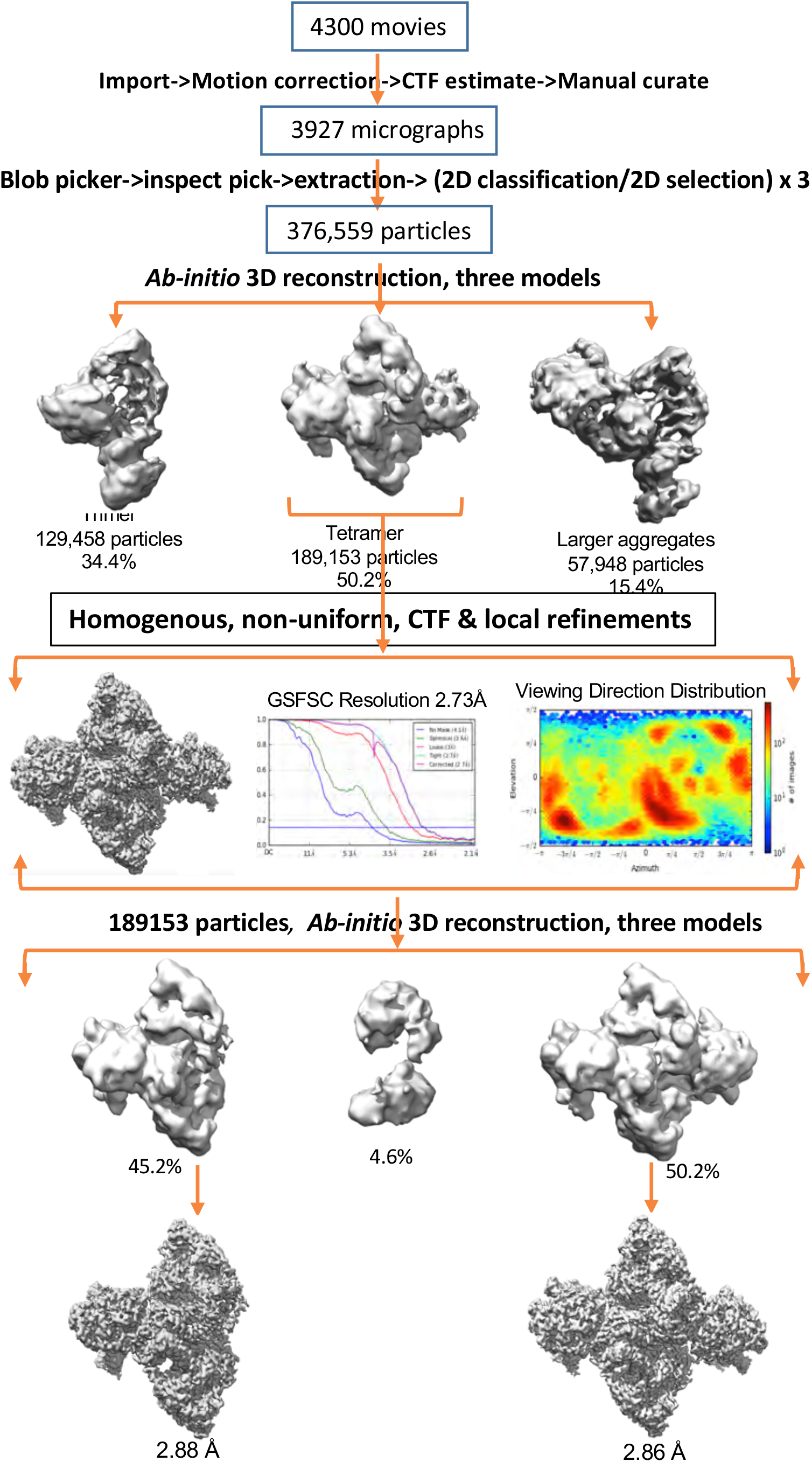
Flow Chart for CryoEM analyses using a KRIOS microscope operating at 300 kV. Computational processing, 3D reconstruction and refinement corresponds to data collected at pixel size of 0.537 Å. For full details of data collection and processing approach, see Methods.

**Supplemental Figure S4.**
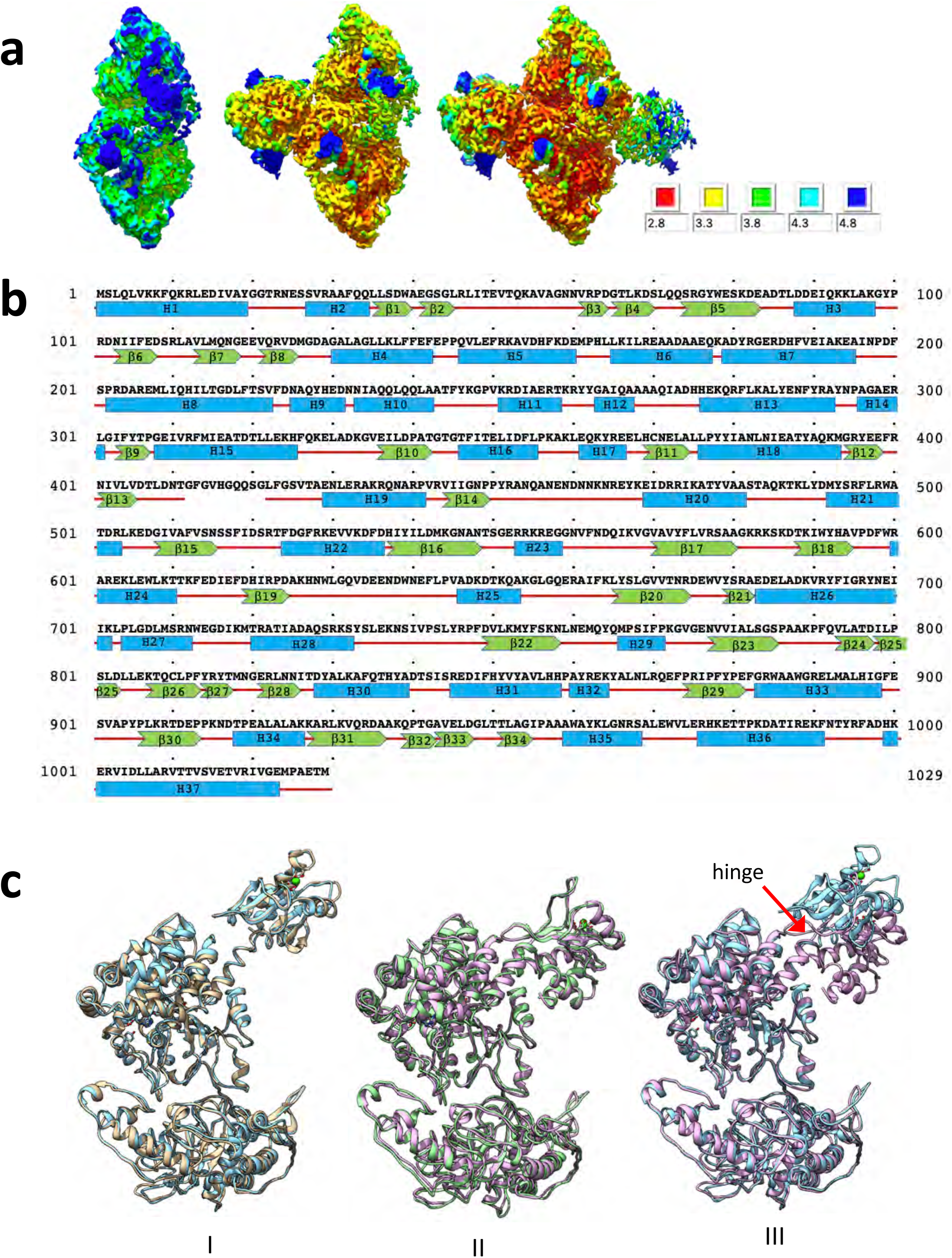
EM resolution and enzyme sequence versus structure. ***Panel a.*** Local resolution distribution of the density maps for the dimer, trimer and tetramer on the same scale. ***Panel b**.* Amino acid sequence and visualized secondary structure of a DrdV subunit. ***Panel c.*** Pairwise secondary structure superposition of the two protomers of the core dimer (I), of the two protomers on the side (II) and one each of the protomer in the center and on the side (III)., showing the conservation of the folding of all domains and their relative disposition in all the particles. The only observable difference in the disposition is a hinged rigid body rotation of the nuclease domain of the protomer on the side with respected to that of the protomer in the core dimers.

**Supplemental Figure S5.**
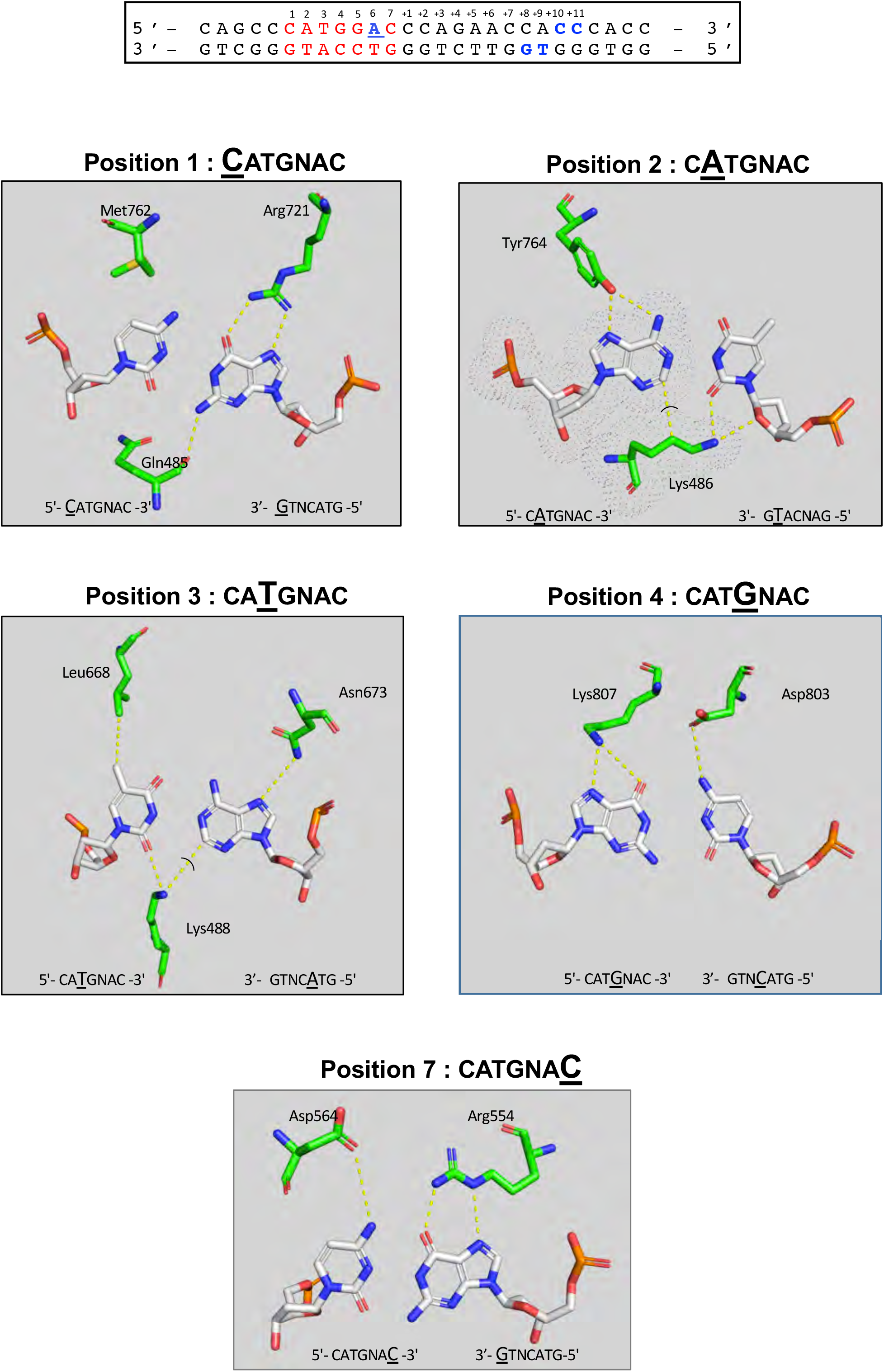
Contacts between DrdV and base pairs in the enzyme’s target site.

**Supplemental Move M1. Rotation and visualization of CryoEM map and corresponding molecular model of the DNA-bound DrdV tetrameric assemblage.**

**Supplemental Move M2. Morph from CryoEM map of DNA-bound DrdV dimeric assemblage to DNA-bound DrdV trimeric assemblage.**

**Supplemental Move M3. Morph from CryoEM map of DNA-bound DrdV trimeric assemblage to DNA-bound DrdV tetrameric assemblage.**

